# Estimating the respective contributions of human and viral genetic variation to HIV control

**DOI:** 10.1101/029017

**Authors:** István Bartha, Paul J McLaren, Chanson Brumme, Richard Harrigan, Amalio Telenti, Jacques Fellay

## Abstract

We evaluated the fraction of variation in HIV-1 set point viral load attributable to viral or human genetic factors by using joint host/pathogen genetic data from 541 HIV infected individuals. We show that viral genetic diversity explains 29% of the variation in viral load while host factors explain 8.4%. Using a joint model including both host and viral effects, we estimate a total of 30% heritability, indicating that most of the host effects are reflected in viral sequence variation.

---

There are differences in the rate of disease progression among individuals infected with HIV. An easy to measure and reliable correlate of disease progression is the mean log viral load (HIV RNA copies per ml of plasma). Measured during the chronic phase of infection, the viral load (referred to as setpoint viral load, spVL) exhibits large variation in a population. Several studies have been carried out to elucidate whether this variation is primarily driven by host genetics ^1–4^, viral genetics ^5–9^, or environmental effects ^7^. Genome-wide association studies consistently show that amino acid polymorphisms in the peptide binding groove of the HLA-A and HLA–B proteins are associated with the viral load of an individual. Furthermore, variants in the HLA-C and CCR5 genes have also been shown to impact spVL. However, those host factors explain less than 15% of the observed phenotypic variance ^3^. In contrast, viral genetic studies and studies of donor-recipient transmission pairs established that 33% of the phenotypic variance is attributable to the transmitted virus itself^5^.

HIV is an extremely variable and adaptive organism with a rapid replication time, and high rates of mutation. Within-host evolution of the viral population occurs during the chronic phase of infection in which the pathogen adapts to its host environment. Several studies showed that a major proportion of the viral sequence is under selective pressure in the host environment, and several viral amino acid changes are associated with host genetic variants in the Human Leukocyte Antigen (HLA) genes ^10,11^.

Viral strains harbor epitope sequences that can be presented by HLA class I proteins of the infected host, which allows the detection and killing of infected cells. The viral population evades detection through escape mutations that modify the epitope sequence but may incur a fitness cost. Compensatory mutations may follow until the viral population reaches its optimal place in a sequence space constrained by the host immune system ^12^.

We collected paired viral/host genotypes along with spVL measurements of chronically infected individuals to estimate the respective contribution of host and viral genetics to the variation in spontaneous HIV control. We used linear mixed models (LMMs, as implemented in the gcta software package ^13^) to explain inter-patient differences in spVL while taking into account host and viral genetic relatedness. LMMs use the pairwise relatedness of individuals with respect to a large set of features (rather than the full individual level data itself) to estimate the fraction of phenotypic variance attributable to those features. Such models have been successfully applied to estimate narrow-sense heritability from genome-wide genotype data ^13^. Concurrently, LMMs were proposed to incorporate phylogenetic relatedness between samples in comparative analyses ^14^, a technique that was further developed to estimate the viral genetic contribution to spVL ^8^. We estimated the respective contributions of host and viral genetics to spVL by defining two relatedness measures, one with respect to the host genotypes, the other with respect to the viral genotypes, and used these jointly in a linear mixed model.

We focused on amino acid variations in the HLA-A, B and C genes due to their established associations with HIV control ^3^. In particular, we used 33 amino acid polymorphisms selected by L1 regularized regression ^15^ to represent the genetic relatedness of the host (**Supplementary Table 1**). Principal component analysis based on host genome-wide genotype data confirmed the lack of major population stratification in the host sample.

The pairwise genetic relatedness of the dominant viral strains observed in the samples was calculated from phylogenetic trees similarly to ^6^. Nucleotide sequences were translated to amino acid sequences, which were in turn aligned with MUSCLE ^16^ and used to derive the correct codon-aware nucleotide alignment. The phylogenetic tree was built from the aligned nucleotide sequences using RAxML ^17^, which were in turn rooted to the HIV M group ancestral sequence; the whole tree was scaled with the inverse of the median height of the branches. The genetic relatedness of two samples in a given phylogenetic tree is the amount of shared ancestry, i.e. the distance from the root of the tree (excluding the outgroup) to their most recent common ancestor ^18^.

Bulk sequences of three viral genes, genome-wide genotypes and viral load data were collected from 541 individuals of Western European ancestry from Switzerland and Canada infected with HIV-1 Subtype B ^10^ (phenotype distributions are shown in **Supplementary Figure 1**). Viral sequences and spVL data were collected at least two years after infection (in the Swiss cohort) or late in infection (in the Canadian cohort) but prior to the initiation of antiretroviral therapy. Thus, the viral genotypes reflect the result of natural adaptation of the pathogen to the host environment. The viral sequences for 1262, 2187, 548 nucleotides of the *gag*, *pol* and *nef* genes were available for at least 80% of samples studied. Overlapping viral genomic regions were excluded from *gag*, to avoid duplicated sequences in the analysis.

We built three LMMs, one containing human variants, one derived from phylogenetic trees, and one including both host and virus information (**Figure 1**). All models included a binary variable indicating the cohort as a fixed effect. Variance components were estimated by residual maximum likelihood. The genetic relatedness matrix created from 33 amino acid polymorphisms of the human class I HLA genes explained 8.4% (SD=4%) of the observed variance in spVL. In contrast, 28.8% (SD=11%) of phenotypic variation was attributable to the viral phylogenetic tree. Combining the two relatedness matrices in one model yielded a total variance explained of 29.9% (SD=12%), less than the sum of the latter two models. Thus, we show that MHC polymorphisms do not explain additional phenotypic variance beyond viral sequence variation.

**Figure 1.**
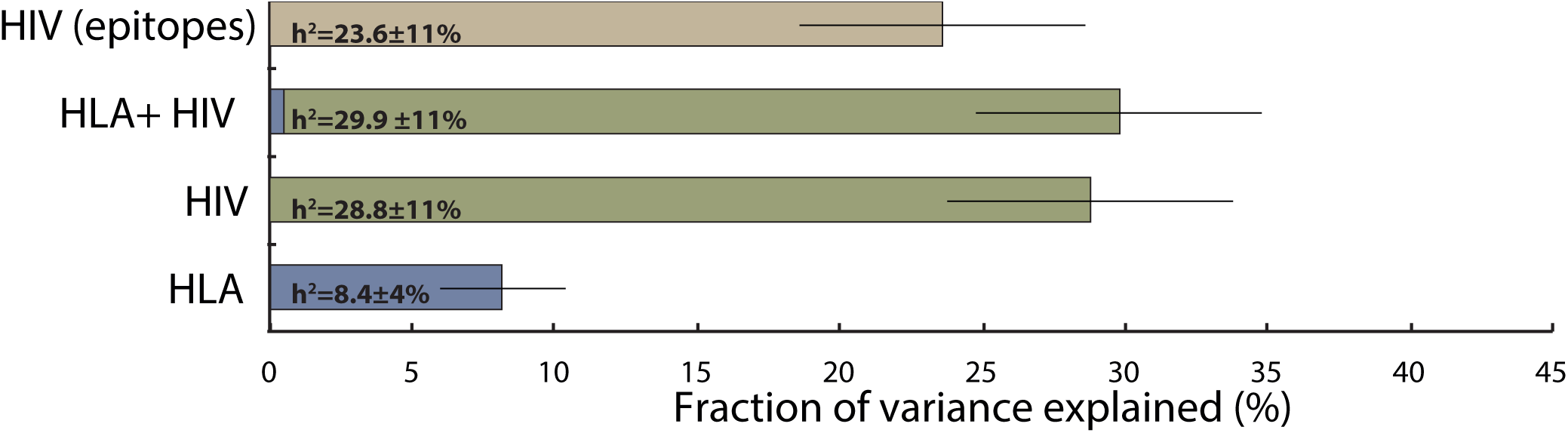
Illustration of fractions of explained variances by models taking human HLA, viral sequence similarities, or both into account.

We next assessed the contribution of viral variants most likely to have an impact on spVL. These included amino acids in known CTL epitopes ^19^ and those positions whose variation is associated with host genetic variation ^10^ (82%, 60% and 84% of *gag*, *pol*, *nef* codons respectively, **Supplementary Table 2**). We used phylogenetic trees built from those codons to show that viral variation in epitopes or other HLA-associated positions explain 23.6% (SD=11%) of phenotypic variance. This suggests that the majority of the impact of viral sequence on spVL is mediated through positions under host selection.

The difference between the variance explained by viral variants in epitope sequences and the variance explained by the MHC polymorphisms themselves may be attributed to two effects. First, selected viral variants might provide a better surrogate of the impact of the host genotype than the imputed host amino acid variants we used. Rare host genetic factors outside MHC (e.g. the *CCR5* deletion), as well as environmental interactions may influence viral fitness and these effects are not accounted for in our estimate of host heritability. Second, the difference could partly be due to the effect of viral variation independent of the current host, including transmitted escape mutations, i.e. viral sequence variation carried over from the previous host, rather than induced by the current host ^20^. It has been shown that reversion of some fitness reducing escape variants is very slow, potentially allowing for a transitory but measurable effect on viral load at the population level ^11^. Transmitted escape mutations allow the study of association of viral variants with spVL independently of their cognate HLA genotype.

The existence of a major fraction of variance explained by variation in viral epitopes further supports the hypothesis that adapted viral variants directly impact spVL, while HLA class 1 variation, at the population level, is the driver of those viral variants. This result also suggests that host genetic association studies not taking the virus into account underestimate the population level effect of host genetic variation. Combining host and pathogen data provides additional insight into the genetic determinants of the clinical outcome of HIV infection, which can serve as a model for other chronic infectious diseases.

---

**Supplementary Figure 1** - Distribution of HIV setpoint viral load values in the Swiss (SHCS) and Canadian (HOMER) cohorts

**Supplementary Table 1** - List of human amino acid variants in HLA-I genes selected by L1 regularized regression and used throughout the paper.

**Supplementary Table 2** - List of MHC-associated HIV amino acid positions based on epitope maps ^19^ and previous association studies ^10^

## References

1. McLaren, P. J. & Carrington, M. Nat. Immunol. 16, 577–583 (2015).

2. McLaren, P. J. & Fellay, J. Curr. Opin. HIV AIDS 10, 110–115 (2015).

3. Pereyra, F. et al. Science (80-. ). (2010). doi:10.1126/science.1195271

4. Fellay, J. et al. PLoS Genet. 5, e1000791 (2009).

5. Fraser, C. et al. Science (80-. ). 343, 1243727–1243727 (2014).

6. Hodcroft, E. et al. PLoS Pathog 10, e1004112 (2014).

7. Mackelprang, R. D. et al. J. Virol. 89, 2104–2111 (2015).

8. Alizon, S. et al. PLoS Pathog. 6, e1001123 (2010).

9. Müller, V., Fraser, C. & Herbeck, J. T. Viruses 3, 204–216 (2011).

10. Bartha, I. et al. Elife 2, e01123 (2013).

11. Kawashima, Y. et al. Nature 458, 641–5 (2009).

12. Schneidewind, A. et al. J. Virol. 82, 5594–5605 (2008).

13. Yang, J., Lee, S. H., Goddard, M. E. & Visscher, P. M. Am. J. Hum. Genet. 88, 76–82 (2011).

14. Housworth, E. a, Martins, E. P. & Lynch, M. Am. Nat. 163, 84–96 (2004).

15. Tibshirani, R. J. R. Stat. Soc. Ser. B 267–288 (1996).

16. Edgar, R. C. Nucleic Acids Res. 32, 1792–7 (2004).

17. Stamatakis, A. Bioinformatics 30, 1312–1313 (2014).

18. Felsenstein, J. Genetics 125, 1–15 (2007).

19. Yusim, K. et al. Los Alamos, NM Los Alamos Natl. Lab. Theor. Biol. Biophys. 3–24 (2009).

20. Van Dorp, C. H., van Boven, M. & de Boer, R. J. PLoS Comput. Biol. 10, e1003899 (2014).

